# Atlas-Based Spatio-temporal MRI Phenotyping of 3D Fungal Spread in Grapevine Wood

**DOI:** 10.64898/2026.01.05.695522

**Authors:** Gargee Phukon, Maïda Cardoso, Christophe Goze-back, Loïc le Cunff, Jean-Luc Verdeil, Cédric Moisy, Romain Fernandez

**Author notes:** Address correspondence to (R.F.).

## Abstract

In perennial crops, inner wood degradation by pathogens often escapes detection until irreversible damage has occurred. Grapevine trunk disease (GTD) is a well-known example in viticulture that alters plants from within, years before foliar symptoms arise, making early assessment difficult. To overcome this limitation, we present a novel non-destructive 3D + t pipeline for Magnetic Resonance Imaging (MRI) spatial quantification and monitoring of early internal tissue degradation resulting from fungal colonization. This pipeline integrates (i) anatomical alignment and rigid time-series registration of volumetric MRI scans, (ii) a generalized cylindrical coordinate transformation for cross-sectional trunk anatomy normalization, (iii) supervised classification to segment water-depleted (diseased/non-functional) regions, and (iv) population-level statistical analyses including construction of population mean images, probabilistic atlases of lesions, and 3D lesion descriptors. Applied to multiple Vitis vinifera cultivars inoculated with a fungal trunk pathogen, our approach enables time-lapse comparisons between cultivar and treatment in vivo. The results reveal consistent early degradation signals across individuals and cultivar-dependent lesion differences. By combining high-resolution MRI with advanced image processing and statistical atlas tools, this method provides a new paradigm for 3D plant phenotyping of internal disease progression. This methodological innovation allows non-invasive quantification of disease development and comparative assessment of host responses in woody plants, demonstrating its potential to advance understanding and management of GTDs.

## INTRODUCTION

Viticulture plays an important role in the cultural and economic heritage of the Mediterranean region and is a major agricultural sector worldwide. Recently, grapevine trunk diseases (GTDs) have become a significant challenge to the long-term sustainability and productivity of vineyards. Common GTDs, such as Eutypa dieback, Esca and Botryosphaeria progressively damage the internal woody tissues of grapevines, leading to declining vine health, reduced yields, and eventually vine death, causing significant economic losses for growers [1]. Detection and monitoring of GTD involve tracking foliar symptoms like leaf discoloration or patterns. They are unreliable because their onset may occur within three to eight years after infection [2]. Hence, while visual scouting remains fundamental, it does not actually describe the internal health state of the specimen.

Traditional methods for controlling GTDs are often destructive, such as trunk removal, or rely on toxic fungicides still used in some countries but now banned in the European Union (EU). In this context, our objective is to develop non-destructive methods to characterize the early progression of GTDs in the internal tissues, quantify the disease development and differentiate early varietal responses to alternative treatment.

Because conventional approaches are destructive or unreliable, non-destructive imaging techniques have been brought forward to overcome this challenge. Over multiple years, ground-based multispectral and hyperspectral imaging has been proposed as an innovative tool to enable automated detection of Esca-symptomatic vines [3]. However, this method is limited to capturing temporary external factors that have been shown to be poorly correlated with the actual health status of the plant and its phytopathological fate. Mid-infrared (MIR) spectroscopy [4] was introduced as a fast, nondestructive alternative to identify GTD pathogens, but the diagnostic accuracy is structurally limited by shallow penetration, sensitivity to sample moisture, and difficulty in distinguishing among different trunk diseases, such as those caused by fungi versus other agents, which have distinct long-term consequences for the vine.

All these approaches mainly focus on what can be observed externally or within a shallow layer of wood, whereas GTDs develop primarily in the deeper structures of the vine. Consequently, X-ray computed tomography (CT) and micro-computed tomography (microCT) [5], [6] emerged as promising tools to study the internal structure of grapevine trunks. CT can reveal differences in radiodensity, and its micro-CT variant delivers exceptional resolution, enabling precise identification of internal defects, including necrosis, decay, and black spots. CT provides a full 3D structural description of grapevine wood [7]; however, it does not capture the plant’s physiology, including responses to pathogens and therefore cannot detect the weak signals associated with early stages of fungal infection, before structural degradation occurs and becomes detectable by CT. This limitation calls for complementary investigation techniques such as magnetic resonance imaging (MRI).

In plant biology, MRI has been used to observe both the static anatomy of internal tissues and dynamic processes like fluid flow. The primary advantage of MRI over CT is its ability to provide details about both the structure and the function of plant tissues. High-resolution 3D MRI images can be leveraged in plant imaging for measuring physiological parameters like water content, flow velocities, and tissue water status (mechanical and chemical water binding). MRI can be tuned to different modalities (e.g. T1-weighted, T2-weighted, diffusion weighted imaging) to highlight different aspects of plant tissues. For instance, T1-weighted images highlight watered tissue boundaries, while T2-weighted images are more sensitive to the water chemical and mechanical binding [7]. Based on these capacities, numerous applications have been developed in plant research, including studying water status of leaves [8], assessing xylem function [9], monitoring root growth and development [10], [11], quantifying water content and storage [12], and investigating stress physiology and drought responses [13].

Because MRI is non-invasive and non-destructive, it enables repeated monitoring of a specimen over time. It provides information on internal tissues without harming the plant, and its high sensitivity to water distribution and dynamics makes it a promising tool to detect physiological state changes and quantify infection responses. Based on these advantages, we hypothesize that MRI is suitable for non-destructive monitoring of internal tissues and quantitative characterization of grapevine cuttings from diverse varietal backgrounds. To test this hypothesis, we used MRI to monitor *in vivo* disease progression within grapevine trunks, generating and analyzing a four-dimensional dataset comprising 3D anatomical and functional information of specimens acquired at multiple time points. Ultimately, these efforts aim to enable early, non-destructive diagnostics and population-level characterization of trunk diseases using MRI. To develop a methodology for early GTD detection and quantitative characterization, we focused on visualizing and measuring pathogen-induced effects on water content and tissue status in xylem vessels, cambium, and rays surrounding inoculation sites. Thus, the objective of this present work is to establish a methodological framework and test its suitability for longitudinal *in vivo* monitoring of disease effects, while assessing differences in susceptibility among grapevine varieties to an Esca-associated pathogen.

We undertook a systematic evaluation of wounded specimens with and without pathogen exposure. To make these comparisons robust across specimens, we developed an automated image alignment pipeline that registers individual MR images into a common reference geometry. Given that grapevine trunks naturally vary in diameter, curvature, and anatomy, direct comparison of infection patterns across specimens is challenging. To address this, we propose a custom transformation framework to resample trunk cross-sections relative to the cambium contour in all the 3D volumes. We designed a generalized cylindrical transform to unwrap the curved cambium into a flat reference line, producing a standardized geometry across specimens. Such a normalized coordinate system is expected to enable radial (Δr) and tangential (Δθ) progression to be clearly separated, thereby highlighting whether pathogen spread occurs along the cambium or into the xylem rays.

We hypothesize that this method can be automated and applied high-throughput to facilitate the quantification and comparison of complex circular infection patterns across populations. However, many studies that use advanced imaging and geometric analysis, including 3D + t geometry, are limited to a single specimen, or only a few individuals. This is often due to the complexity of experimental protocols, the challenges of large-scale data management, and the difficulty of comparing specimens that vary in size, shape, or geometry. As a result, most imaging-based studies lack statistical power and cannot generalize their conclusions to a population. Interestingly, population averaged images have been long used in atlas-building frameworks in biomedical imaging, (such as the ICBM/MNI brain atlas [14]) and atlas frameworks became instrumental in computational anatomy. Recent imaging studies consistently rely on mean images to represent population-level signal patterns, including fetal [15] and neonatal MRI [16], and population based organ atlases [17]. However, the application of atlases to plant imaging is relatively new [18].

In this study, we present a methodological framework combining 3D + t MRI, geometric normalisation, and probabilistic atlas construction to enable population-level analysis of early grapevine trunk disease progression. As a proof of concept, we applied this pipeline to grapevine cuttings inoculated with *Phaeoacremonium chlamydospora* and monitored lesion development over time across multiple cultivars. Using atlas-based representations and geometric lesion descriptors, we show that early internal tissue degradation can be quantified reproducibly and compared across individuals. Notably, these measurements reveal distinct early response patterns among cultivars, allowing the identification of response groups at early stages of infection. Interestingly, the susceptibility ranking inferred from early MRI-derived lesion dynamics does not fully match the long-term classifications reported in the literature, raising new questions about the sequence linking early physiological responses and chronic trunk disease symptoms.

## MATERIALS AND METHODS

**Fig. 1:**
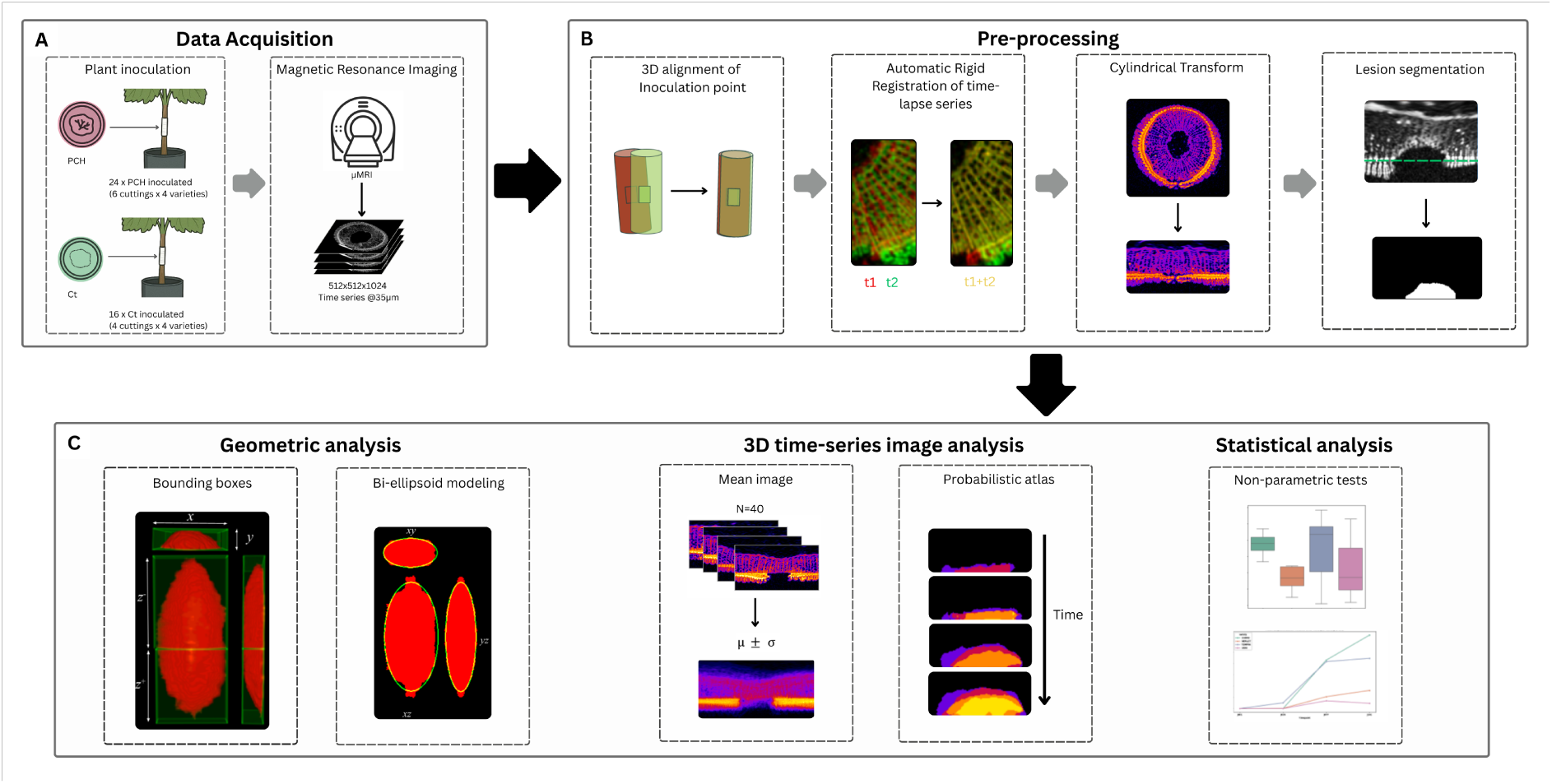
Schematic overview of the methodological pipeline. (A) Plant material was inoculated with the fungal pathogen or control and imaged at successive timepoints using µMRI. (B) The resulting 3D images were pre-processed through alignment of the inoculation point, automatic rigid registration of time-lapse datasets, cylindrical transformation of the trunk volume, and lesion segmentation. (C) Quantitative analysis was then performed using geometric lesion descriptors, including bounding box and bi-ellipsoid modeling. Group-level comparisons were then carried out using mean images, probabilistic atlases, and statistical comparisons to assess lesion development across conditions and varieties.

### Plant and fungal material

The experiment was conducted on 40 grapevine stem cuttings (ten repetitions for each of four cultivars). Each cutting was approximately 15 cm in length, with two buds. After pruning, cuttings were individually potted in plastic tubes designed to fit the micro-MRI chamber (40 mm diameter). The cuttings were maintained in a greenhouse at CIRAD, France under controlled conditions, including regulated temperature and humidity, supplementary lighting, automated irrigation, and preventive treatments against downy and powdery mildew.

#### Vine cultivars

We selected four cultivars of *Vitis vinifera*, two red (Merlot, Tempranillo) and two white (Chardonnay, Ugni Blanc), representing contrasted responses to GTDs, based on external foliar symptom expression (see **Table 1**). These varieties are major European wine cultivars and differ in their susceptibility to Esca and its associated pathogen *Phaeomoniella chlamydospora* (Pch). By including cultivars considered relatively tolerant (Merlot, Chardonnay) or more susceptible (Tempranillo, Ugni Blanc), we designed the experiment to quantify various responses under infected and uninfected conditions.

**Table 1:**
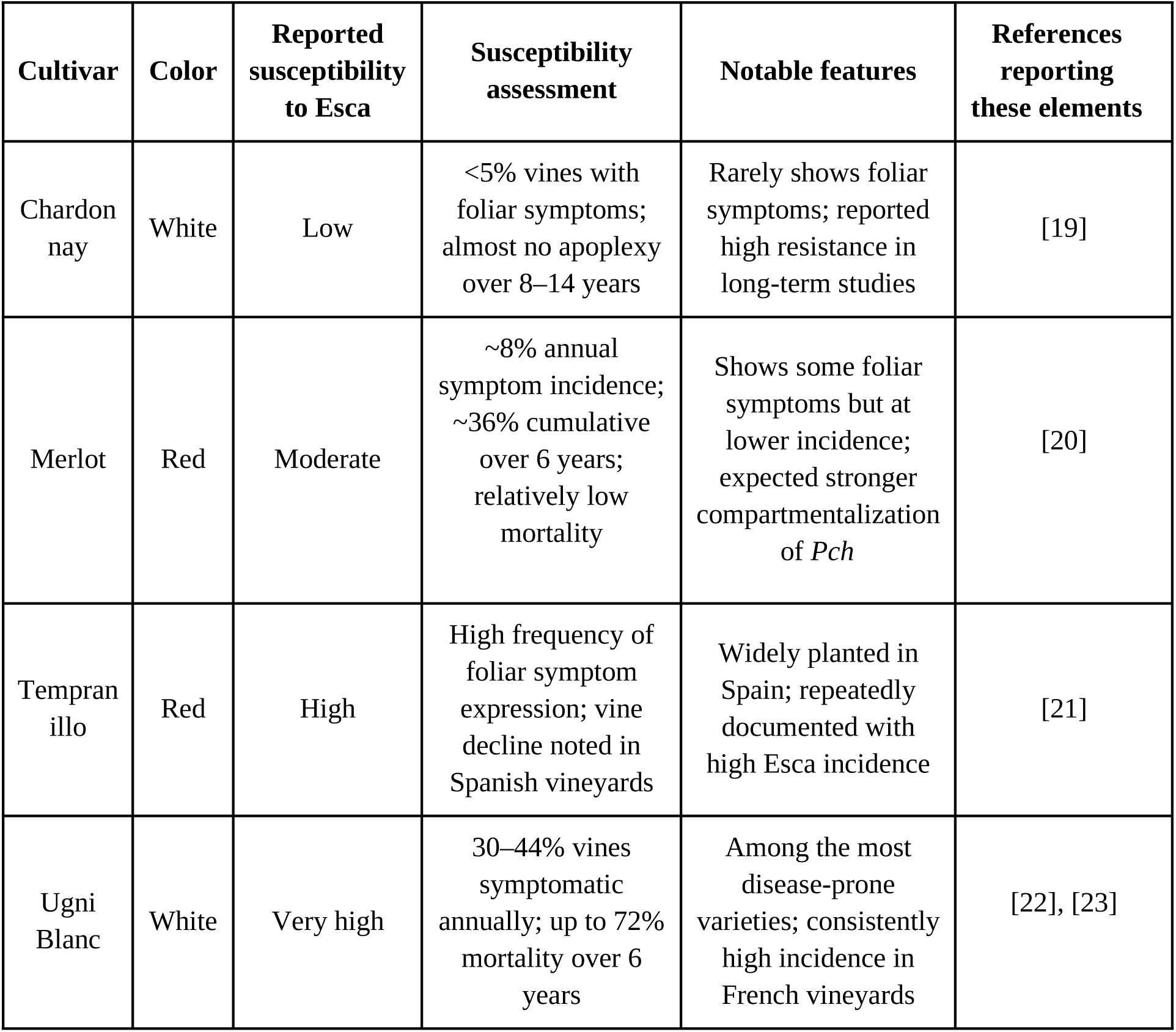
Grapevine cultivars selected for this study and their reported susceptibility to Esca.

While these descriptions cover long-term impact of infection on grapevines, in contrast, our experiment does not focus on chronic outcomes (or year-to-year symptom expression), but instead targets the early stages of fungal infection, caused by artificial inoculation of Pch. This raises a key question of whether the susceptibility gradient that becomes evident over years in the vineyard (especially with Chardonnay generally more tolerant and Ugni Blanc highly sensitive) translates into similar trends in the earliest physiological changes revealed by MRI.

#### Fungal material

*Phaeomoniella chlamydospora* (Pch) is a vascular ascomycete fungus widespread in vineyards worldwide. It colonizes the xylem of grapevines [24], causing vascular damage that is characteristic of Petri disease, and constitutes a core component of the Esca complex in mature vines. Pch spreads through pruning wounds or graft unions, leading to vessel blockage, wood necrosis, and occasionally foliar symptoms or vine death. Unlike wood-decaying fungi, it does not degrade lignin or cellulose but impairs vascular function via toxin production and vessel occlusion. While host defenses such as phenolic accumulation and PR protein expression occur, they are often too delayed to prevent disease. Pch spreads both longitudinally through xylem vessels and radially/tangentially through tracheary tissues.

### Wounding and Inoculation

A standardized square wound of 2 mm × 2 mm was made on each cutting by removing the bark. Over ten wounded cuttings per cultivar (repetitions), six cuttings were inoculated with *Phaeomoniella chlamydospora* (Pch), which was cultured in the laboratory, while the other four received a control treatment. The inoculations were performed as described in [25]. Control treatments were performed by introducing sterile agar plugs into the inoculation sites and resealing them in the same manner as the fungal inoculated specimens. This ensured that control vines experienced the same wounding and handling as inoculated vines, but without the presence of fungal pathogens. This experimental design allowed us to test the hypothesis that Pch-inoculated cuttings would display distinct internal symptom development, whereas control cuttings would primarily undergo normal wound healing, providing a clear contrast between pathogenic progression and tissue response to the wound.

### Time-lapse Magnetic Resonance Imaging

Once the cuttings were inoculated with Pch or control treatment, they were transferred to BioNano Imaging Foundry (University of Montpellier) for magnetic resonance imaging (MRI) (Fig. 2 (C)). Imaging was performed at four time points post-inoculation. The first scan (day 1 post-inoculation) served as a reference image, before visible symptoms development. The second scan (day 29) was expected to capture the early response phase, when localized functional changes associated to water availability in the tissues would begin to appear around the inoculation site. The third scan taken on day 77 post-inoculation was intended to monitor the mid-progression phase, a stage where more extensive tissue alterations were anticipated. The final scan on day 141 was chosen to represent the late stage of disease development, when the most pronounced internal alterations were expected. Collectively, these four time points yielded a time-lapse volumetric dataset (3D + t), allowing spatio-temporal analysis of both structural and functional changes within the same cuttings. The imaging was conducted between June and December 2024.

**Fig. 2:**
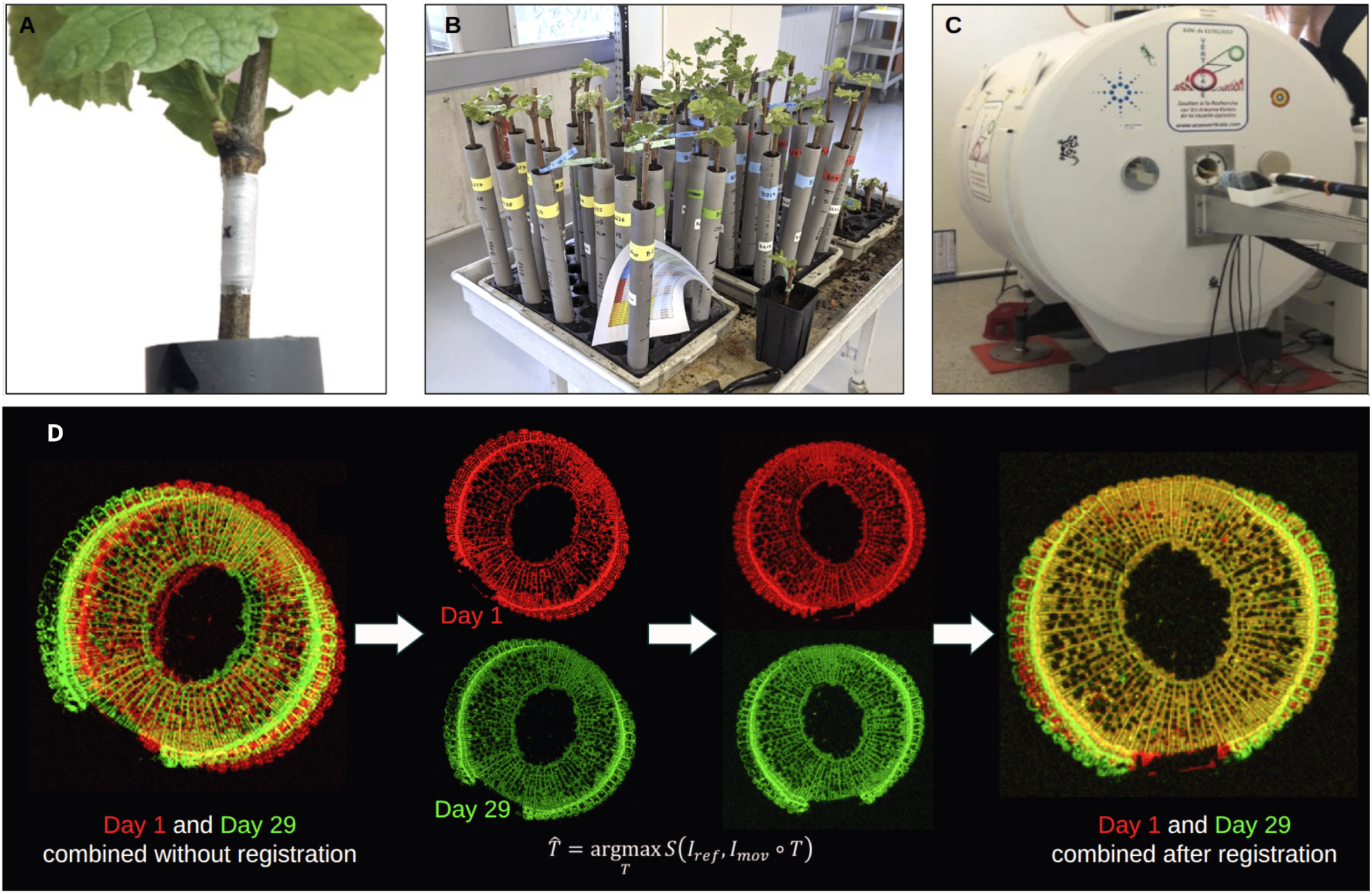
Experimental material, imaging setup, and effect of rigid registration on µMRI data. (A) Grapevine cutting showing the inoculation site after treatment (B) Total population stored in the greenhouse (C) MRI at BioNanoNMRI, Montpellier University (D) Representative transverse µMRI slices acquired at two timepoints (Day 1 and Day 29), shown before and after rigid registration. Overlaying images without registration highlights spatial mismatches, while rigid registration aligns internal structures, enabling voxel-wise comparison of tissue changes over time.

All MRI acquisitions were performed using a 9.4 Tesla high-field MRI system with 35 µm isotropic resolution. This high-resolution scanner enables detailed anatomical visualization of wood integrity and tissue degradation. Three MRI modalities were employed: Ge3D, T1-weighted and T2-weighted. In this study, we used only the Ge3D data, that are high-resolution and isotropic, in order to foster an accurate geometrical description of the phenomenon.

### Pre-processing

Each of the 160 resulting 3D volumes (one per specimen per time point) contains over 1,000 serial slices with a 512x512 definition. The images are 32-bit multi-slice TIFF files (1GB per stack). Within these 3D volumes, the specimen appears with random orientations and inconsistent positions within the image coordinate system (x,y,z). To enable meaningful comparisons, a multi-step processing pipeline [26] was designed, consisting of normalization, registration, cylindrical transformation and segmentation.

#### Normalization

To ensure consistent comparison of image intensities across specimens and time points, all MRI stacks were normalized using a water-filled capillary included in every scan as an internal reference. Voxel intensities were linearly rescaled relative to the median reference signal after subtracting background noise. This produced standardized image stacks where values are expressed in relative units with respect to the capillary, thereby reducing the influence of the device variability and enabling quantitative comparisons across time and treatments.

#### Inoculation Point Alignment

Raw images differed in placement because samples were mounted manually, meaning the specimen would appear at slightly different positions and orientations across datasets and time series. The alignment step was intended to correct these inconsistencies by reorienting and repositioning each 3D image stack so that all specimens shared the same reference geometry. By treating the inoculation point as an invariant reference, we eliminated misleading sources of geometric variability across datasets and established a stable ground for analyzing patterns of growth and the effects of infection spread over populations.

From each grapevine MRI scan, we manually extracted the real-world coordinates of three anatomical landmarks, the approximate midpoints of the trunk in the top and bottom slices, and the centre of the inoculation site. From these points, an orthonormal basis was constructed. The vector from the bottom to the top landmark defined the longitudinal axis of the trunk, ***v_z_***. The inoculation point was then orthogonally projected onto this axis, allowing its position to be expressed in two parts: an axial component describing inoculation point height along the trunk, and a lateral component describing the depth in the trunk. The lateral offset vector defined the second axis, ***v_y_***, while the third axis ***v_x,_*** was obtained by calculating the cross product, ensuring an orthogonal right-handed reference frame. Using this basis, a linear transformation was computed that reoriented the specimen into the desired axes and translated the inoculation point to a predefined target coordinate in voxel space.

#### Rigid Registration

We used automatic rigid registration with Block-Matching from the Fijiyama libraries (Fernandez et al., 2021) to align specimens across different timepoints. While MRI signal intensities change during fungal progression, the wood tissue geometry remains stable over the time interval of the experiment. The biologically relevant variability occurs primarily within internal tissues, which can be meaningfully tracked once images are aligned in a common spatial framework. To that end, we used rigid transformation that includes only translation and rotation while preserving distances, angles, and anatomical proportions. As a result, the shape of the wood and its internal anatomical structures remain geometrically unchanged by the registration process, while the intensity of the corresponding tissues could change with time, during the ongoing progression of the pathogen, highlighting change in watering status, and allowing an accurate identification of pathogen progression.

Rigid registration (Fig. 2(D)) was performed in a daisy-chained manner to align all 3D volumes to Day 1, which was used as the reference scan. Day 29 was registered directly to Day 1 (T2→1), together with the inoculation alignment. Day 77 was first aligned to Day 29 (T3→2) and then mapped to Day 1 by composing T3→2 with T2→1, together with the inoculation alignment. Similarly, Day 141 was registered to Day 77 (T4→3) and brought into the Day 1 space via application of the composed transformation T4→3, T3→2, T2→1, and the inoculation alignment.

Following registration, each 3D image volume was stacked into a 4D hyperstack, where the Z-dimension represents the image depth (slices), and the T-dimension corresponds to the time points (Days 1, 29, 77, 141 post-inoculation). This final hyperstack enables synchronized temporal and spatial analysis across the different modalities and timepoints for each specimen.

#### Generalized Cylindrical Transform

Woody perennials present considerable heterogeneity in size, shape, and tissue composition, making direct comparison across specimens challenging. However, grapevine cuttings exhibit distinctive elements of similarity: they have a cylindrical form and their cross-sections are almost a circle, that do not change within the experiment duration (the wood keeps its shape, even infected). To facilitate comparison across samples of different shapes and sizes, we unwrap these circular cross-sections into a flat representation using a cylindrical transform (2D polar transform along the Z axis). This transform has been widely used across disciplines whenever the underlying structure is circular or cylindrical, from unwrapping retinal [27] and endoscopic images [28], to recognizing contactless fingerprints [29]. By converting our (𝑥,𝑦,*z*) 3D images to cylindrical space (r,𝜃,*z*), we hypothesize that the complex circular progression patterns of the fungus around the cambium and into the xylem become easier to quantify (see Figure 3). In this framework, radial direction (in depth, towards stem center) and tangential directions (along the stem axis or along the cross-section contour) are defined in a simplified manner, providing a basis for analyzing potential effects of infection spread along the denser wood of xylem rays or across it.

**Fig. 3:**
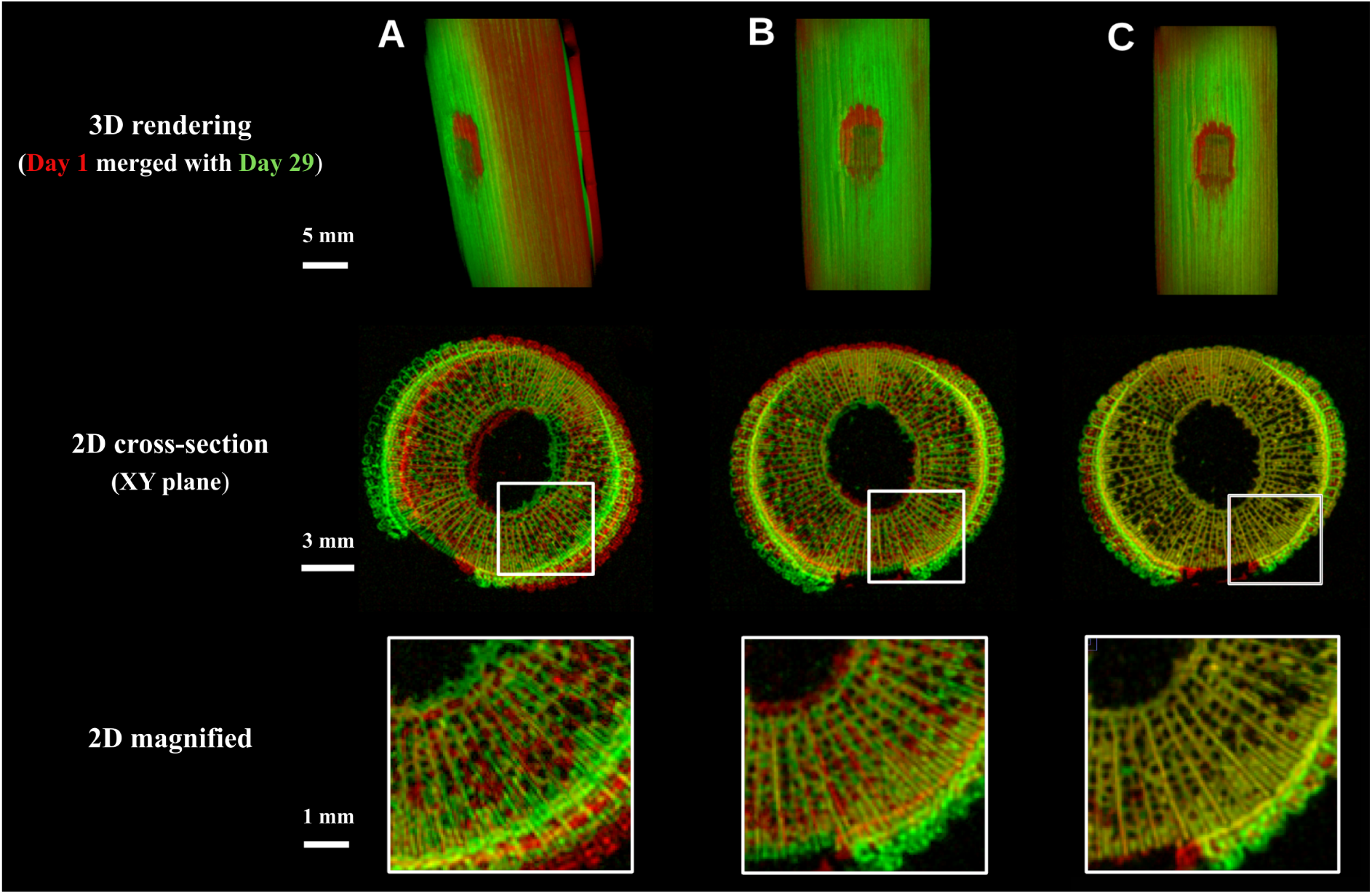
Results of geometric alignment and rigid registration on longitudinal µMRI data superposition. Superimposed µMRI data are shown (A) before processing, (B) after geometric alignment of the inoculation point to a common reference position, and (C) after automatic rigid registration. For each condition, the top row shows a 3D rendering of the trunk, the middle row shows a representative transverse cross-section (XY plane), and the bottom row shows a magnified view of the highlighted region. Improved spatial correspondence of internal structures and lesion boundaries is visible after rigid registration (pixels in yellow combine red (day1) and green (day 29), highlighting a good level of image correspondence). Scale bars: 5 mm (3D rendering), 3 mm (cross-sections), and 1 mm (magnified views).

However, the cross-sections of grapevine cuttings are not perfect circles. These non-uniformities introduce inconsistencies that a standard polar transform cannot handle. To address this limitation, we developed a ‘generalized’ cylindrical transform algorithm that aims to adapt to irregular geometry and produce more consistent and comparable maps of fungal progression. The sequence of 3D images was transformed from the Cartesian space to cylindrical space by resampling along the contour line as it appears on the first time point (prior to any tissue intensity changes caused by the pathogen progression). This transformation unwrapped the cylindrical geometry of the specimen into a flat representation, making the circular cambium appear as a straight line, and disentangling tangential and radial directions. This flattening step was also intended to reduce noise and increase the effective comparability of samples, preserving more of the true signal needed to distinguish subtle differences in fungal behavior between varieties. In Cartesian space, size differences introduced high voxel-wise variance in the population of images when computing mean image, thereby masking biological significance. In contrast, the cylindrical transformation is designed to normalize the geometry, producing more consistent data (area of interest, including wound and lesion progression, share a same geometrical support) in which variability across specimens becomes more informative about the response to the fungal pathogen, and less about the different shapes of the cuttings.

The transformation was performed by sampling points along the cambium of the grapevine trunk with unit spacing between two successive resampling planes. Each resampling plane was defined with its origin on the contour and oriented towards the trunk’s central axis, such that the successive planes intersected at the axis. As the trunk surface is not perfectly circular, the tangential direction is not strictly orthogonal to the radial plane. Consequently, the target pixels in polar space do not correspond to uniform square pixels. Pixels near the cambium are mapped with minimal distortion, whereas pixels closer to the center correspond to smaller circumferences and therefore appear stretched in polar space, forming trapezoidal regions. To compensate for this distortion, we compute a “shrinkage map” (using a *sinθ* factor) which stores, for each pixel, the appropriate surface-area factor required to recover its true physical size. This correction is essential for ensuring that subsequent surface or density computations accurately reflect the underlying tissue geometry.

#### Segmentation

The fungal colonization front cannot be directly observed in the MRI images. However, early fungal activity causes local tissue degradation that results in a loss of MRI signal, reflecting the disappearance of water within the wood and a loss of tissue functionality. This signal loss defines a lesion developing around the inoculation point. We therefore target to segment this lesion, which is a dark, low-intensity area replacing the bright signal associated with healthy, water-rich wood.

As a first step, a machine learning based segmentation model was trained to distinguish watered regions (bright) from non-watered (dark) regions, including dry tissue and background. To train the model, a representative subset of the dataset was assembled. Because lesion development is localized around the inoculation site, only a subset of slices surrounding this region was retained for training. For each treatment and time point, the selected slices from all specimens were gathered and served as training data for a Fast Random Forest classifier using the Trainable Weka Segmentation [30] libraries in ImageJ. The trained model was then applied to the complete dataset to generate binary lesion masks for all specimens across varieties, treatments, and time points.

Due to the pixel-based classification approach of the Trainable Weka Segmentation, low-intensity tissue pixels were often misclassified as non-tissue. To overcome this limitation and obtain the desired masks, we applied a post-processing step based on slice-wise top-fill correction. This approach is straightforward in our setup because, after the cylindrical transform, the upper rows of each slice consistently correspond to tissue, whereas the lower rows represent background. The correction involved identifying the first row with complete (or sufficiently high) tissue coverage and flood-filling all rows above it, ensuring a consistent tissue region. This resulting segmentation provides the basis for subsequent analyses of lesion geometry and progression.

### Quantitative Image Analysis

The previous preprocessing steps were intended to provide a common geometric framework and a consistent localization of the disappearing region, the lesion, across all specimens. Building on this standardized representation, we developed quantitative tools to analyze how the phenomenon varies within and between populations of specimens, particularly between Control and Pch groups and across the different grapevine varieties. To achieve this, we developed aggregation strategies based on mean images and probabilistic atlases, which summarized group-level spatial patterns of lesion growth. We then defined a feature space to describe the 3D + t objects to characterize each specimen with a small number of interpretable parameters. These features formed the basis for the statistical comparisons performed later in the study.

#### Population-Based Atlas Construction

As stated in the introduction, atlas-based approaches are widely used in medical imaging and developmental biology to compare, combine, and quantify complex anatomical structures in 2D, 3D, or 3D + t. These strategies provide population-level descriptions that capture both common patterns and individual variability. In this work, after establishing a common geometric reference frame, we adopted an atlas-based methodology to achieve population-level anatomical phenotyping of the growing lesion.

#### Mean image

Mean images (group-average images) were computed by voxel-wise averaging of the aligned and transformed MRI volumes to obtain a representative signal distribution for each group. Let *I _k_* ( *v*) denote the intensity of the specimen *k* at voxel *v*, with *k* = 1 , … *, N_specimens_* (*I* : ℝ ³ *→* ℝ associating an intensity value to each voxel *v*). then the mean image *μ* (*v*) can be defined as:

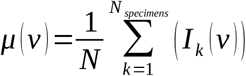

This averaged image captures the first-order trend of the population, reducing specimen-specific noise while preserving the recurrent patterns. Standard deviation maps were also computed to quantify variability around the mean and complement the mean image by highlighting regions where specimens differ.

#### Probabilistic atlas

We constructed a probabilistic atlas that estimates, for each voxel, the likelihood of observing a given tissue state across a group of specimens. The atlas is generated by computing the voxel-wise frequency of lesion occurrence normalized by the number of specimens. We visually assessed the resulting atlas by examining XY and XZ cross-sections and the resulting mean image by analysing the distribution of the MRI intensity values and spatial patterns across individuals.

#### 3D + t specimen-wise geometric feature extraction

While atlas-based representations captured population-level trends, they do not directly quantify the geometric properties of individual specimens to compute statistical comparisons based on these values. To enable geometric comparisons between groups, we extracted a specimen-wise set of interpretable 3D + t geometric features from the segmented lesions. These features helped quantify how the affected wood volume/surface evolves spatially, as the deteriorated region is expected to enlarge over time. From each binary mask, we computed its bounding box to capture the overall lesion extent and we fit an equivalent ellipsoid to the mask to characterize its 3D shape and anisotropy. These 3D + t parameters were then used for characterizing differences between populations, allowing downstream comparisons of lesion progression between Control and Pch treatments and across varieties, at both the inter-variety and intra-variety levels.

#### Specimen lesion bounding box

We defined axis-aligned 3D bounding boxes around the lesion mask volumes to characterize them with a few parameters. The separation of the lesion volume into two halves was necessary because the progression of water disappearance tends to spread unevenly across the Z directions, the impact being often bigger on the upper tissues. To that end, these masks were divided at the centralmost Z slice (z= *cz*) to separate the volume into two complementary halves. The Z^-^ region, comprises slices below the inoculation plane (*z < cz*), and the Z^+^ region comprises the slices above it *(z ≥ cz).* This convention follows the standard geometric notation where the negative and positive indices represent opposite sides of a reference plane.

For each half-volume, an axis-aligned bounding box was computed by scanning the non-zero voxels of the segmented lesion binary mask to determine the minimum and maximum coordinates along *x*, *y* and *z*. Specifically, *x_min_* and *x_max_* correspond to the leftmost and rightmost lesion voxels, *y_max_* to the highest extents in the vertical direction (cross-section-wise), and *z_min_* and *z_max_* to the first and last slices containing lesion voxels.

#### Specimen lesion equivalent ellipsoid

In addition to the bounding-boxes, we aim to describe the lesion with geometrical features that better reflect its actual shape. For this purpose, the segmented lesion volumes were modeled by fitting 3D ellipsoids defined by the parameters *(x_0_,y_0_,z_0_,r_x_,r_y_,r_z_*), because the lesion volumes tend to exhibit such geometry. As the inoculation point lies on the outer surface of the wood, the lesion progresses only towards the centre and not outwards, resulting in an asymmetrical geometry, forcing us to estimate “half-ellipsoids”. Moreover, the lesion may develop differently above and below the inoculation point along the longitudinal axis of the cutting, which motivated the use of a bi-ellipsoid model (an ellipsoid to represent the upper half of the lesion and an ellipsoid to represent the lower half).

To implement this, each half (Z^+^ and Z^-^) of the lesion mask was mirrored along the *y* and *z* axes, producing double-symmetric volumes that approximated ellipsoidal shapes suitable for further geometric fitting. To obtain the desired equivalent ellipsoid representation of each double-symmetric lesion volume, we first extracted the contours of the lesion mask volume and retrieved the corresponding surface voxels, that is the point set ({*x_i_, y_i_, z_i_*)}*^n^_i_*_=1_ defined as the coordinates of lesion voxels that neighboured at least one non-lesion voxel in the 6-connected directions. As the lesion is mirrored, the volume exhibits symmetry across both the *y* and *z* axes, cancelling out the cross-product moments *(xy,yz,xz)*, resulting in a diagonal inertia matrix with no rotation terms. We determine the ellipsoid parameters solving a least-squares problem using singular value decomposition (SVD) that identifies the set of parameters that minimizes the overall fitting error, yielding the ellipsoid centre *(x_0_,y_0_,z_0_)* and three radii *(r_x_,r_y_,r_z_).* By construction*, y_0_* and *z_0_* are the same for all ellipsoids, due to the double mirroring around the inoculation point. The radii described the spatial extent of the lesion along each axis, providing a quantitative measure of how far the infection had spread in different anatomical directions. As a final step, we evaluated the fit quality using residual metrics to ensure the fitted ellipsoids provided an adequate geometric approximation.

## RESULTS

The results are presented in two parts. First, we report the methodological results, evaluating whether each part of the pipeline performs as expected. Second, we apply the validated pipeline to characterize fungal progression within cuttings and quantify the differences between treatment groups and cultivars.

### Methodological Results

This section presents the methodological outputs of the pipeline, including alignment to the inoculation point using geometric references, rigid-body registration, cylindrical transformation, segmentation of the disappearing area, and the construction of population-level models such as mean images and probabilistic atlases. We also report the geometric descriptors extracted from each lesion, namely bounding boxes and equivalent ellipsoids, which are later used for quantitative comparisons between groups.

#### Inoculation point alignment

Manual alignment was based on three landmarks indicating the inoculation point and stem axis. We evaluated the effectiveness of this alignment by calculating the distance between corresponding anatomical points in scans of the same specimen over time. To that end, we used the Evaluate Mismatch functions in the ImageJ Fijiyama plugin [31]. The mean displacement between matched points after this first registration step was approximately 3 pixels in X, 7 px in Y and 0.2 px in Z, indicating that most residual variability occurred in the section plane rather than along the stem axis. In real-world units, these offsets correspond to roughly 0.10 mm, 0.25 mm, and 0.007 mm, respectively. Compared to the inoculation wound size (2mm x 2mm), these offsets are really small. The largest mismatch (Y) represents 12% of the wound size while the axial offset is negligible (<1%). When compared to the average cutting diameter (15 mm), the same mismatches represent <2% of the stems diameter, showing that the global orientation is preserved at the specimen scale. The overall 3D displacement averaged ∼0.27 mm which is equivalent to about 8 voxels given our isotropic voxel size of 0.035mm. This indicates the inoculation-based alignment standardized orientation but left residual offsets that limit precise voxel-wise comparison.

#### Automatic rigid registration

After applying rigid registration, mean displacement was reduced to 0.30 px in X and Y and 0 px in Z, corresponding to a maximum of ∼0.01 mm in each direction, and the total 3D distance dropped to ∼0.02 mm, or approximately half a voxel. Rigid registration improved alignment accuracy from ∼8 voxels to <1 voxel, achieving sub-voxel correspondence of internal anatomical structures. These results suggest that this hierarchical registration strategy combining alignment and automatic block-matching produces a sufficiently consistent framework to support voxel-wise anatomical comparison.

### Generalized Cylindrical Transformation

By construction, the cylindrical transformation performs the “flattening” of the cambium tissue, from a cylinder to a flat plane. However, one limitation lies in the fact that the cross-sectional profile of the resampling cylinder is extracted manually from the central slice at the wound center level. As the object contour is not perfectly invariant along the Z axis, at some distance of the wound the contour shape is slightly changing. As a consequence, after the transformation the cambium appears less flat at a distance from the wound (Fig. 4 (B), red arrow). However, this concern appeared to be unimportant as visual investigations showed that the distorsion is very limited (Fig. 4 (B), green arrows) in the area under study (direct surroundings of the wound).

**Fig. 4:**
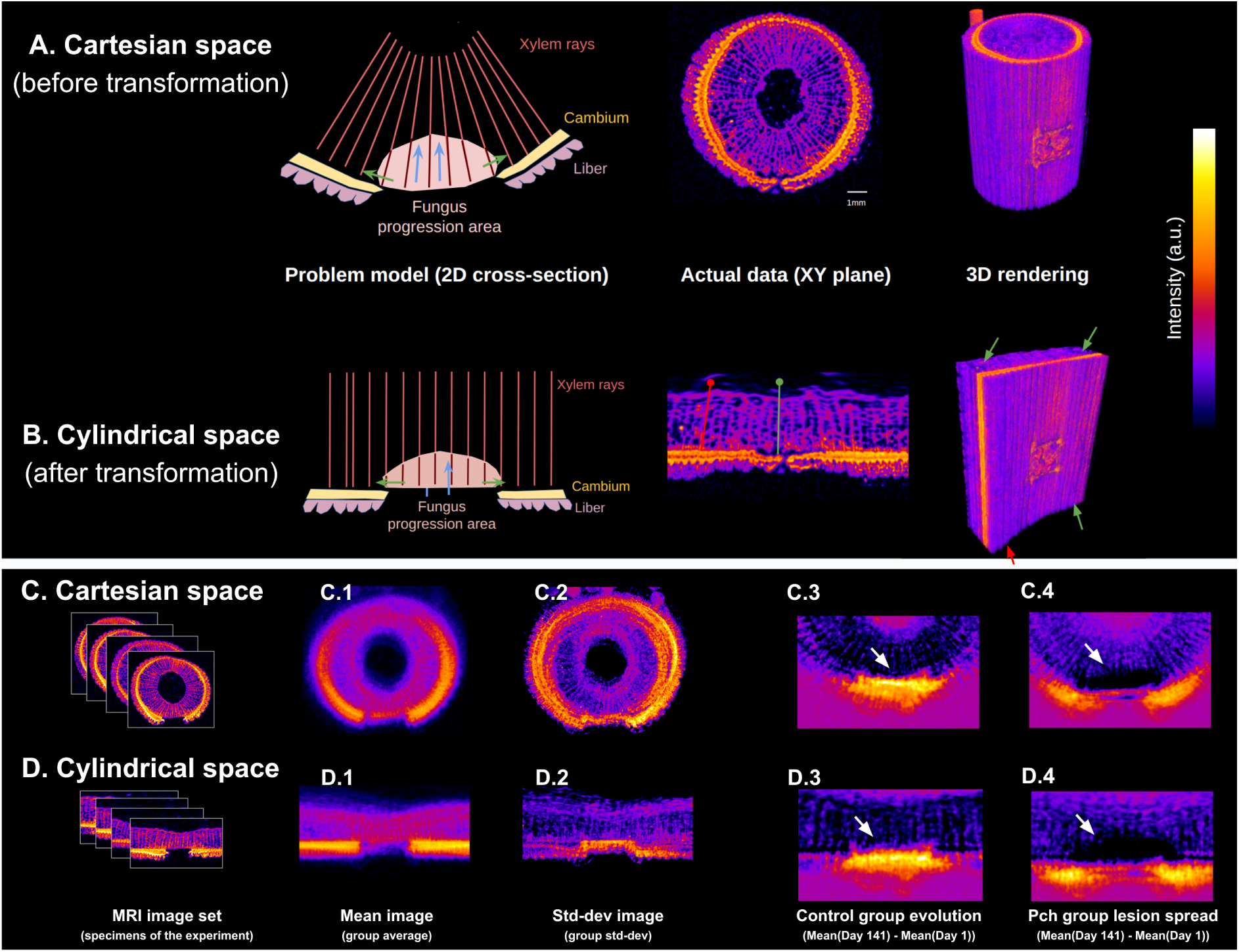
Comparison of Cartesian and cylindrical representations. (A) Illustration of the problem in Cartesian space, showing how fungal progression relative to anatomical structures (xylem rays, cambium, liber) is distorted by specimen curvature, together with representative MRI data displayed as a transverse cross-section (XY plane) and a 3D rendering. (B) Corresponding representation after cylindrical transformation, in which the trunk geometry is flattened into a common spatial framework, enabling consistent localization of the wound and lesion region. (C) Analysis performed in Cartesian space, including mean images (C.1), the standard deviation map (C.2), the difference map in Control (C.3) and Pch (C.4). (D) Corresponding analysis in cylindrical space, showing improved spatial alignment and reduced geometric variability, resulting in clearer mean patterns (D.1), more interpretable variability maps (D.2), and the difference images (D.3 and D.4).

We investigated the quality of separation between radial and tangential axes by measuring the mean deviation of the orientation of xylem rays relative to the expected orientation (vertical), on a random set of 10 images from the four varieties (28 rays measured for each image). After transformation, we measure an almost vertical mean orientation (89.1°), and a 3.07° mean deviation from the vertical axis (see Supplementary materials S.1). These results indicate an effective and unbiased reorientation of rays, while the separation of tangential and radial directions is imperfect, suggesting that the generalized cylindrical model covers most of the cutting geometry, but not totally.

Conceptually, the transformation worked as intended. However, because the fungus remained highly localized, the geometric changes introduced by the transform were limited. Still, this approach could become more informative in longer monitoring setups, or with other pathogens or hosts that show different spread behaviors.

### Segmented Masks

The segmentation step produced binary masks of the disappearing regions across all specimens, treatments and time points. The classifier separated bright (water) tissues from dark (non-water) tissues while the top-fill correction recovered tissue pixels that were initially misclassified as background, restoring a 3D tissue region across slices that appeared to be continuous and anatomically consistent in our visual verifications (see in Fig. 5 (A)). The resulting masks were then inverted and axially cropped around the inoculation point to isolate the region of interest along the stem. These final lesion masks delineated the zones of signal loss, which then served as inputs for atlas construction and bounding box extraction. The boundary pixels of these lesion masks were then extracted as surface points for equivalent ellipsoid fitting. The quality of the masks was verified by merging each segmentation with the original image and visualizing the overlap. In infrequent cases (3 images over 160), segmentation exhibited large discrepancy with the actual contour of the lesion area, and the slice-wise correction was done manually.

**Fig. 5:**
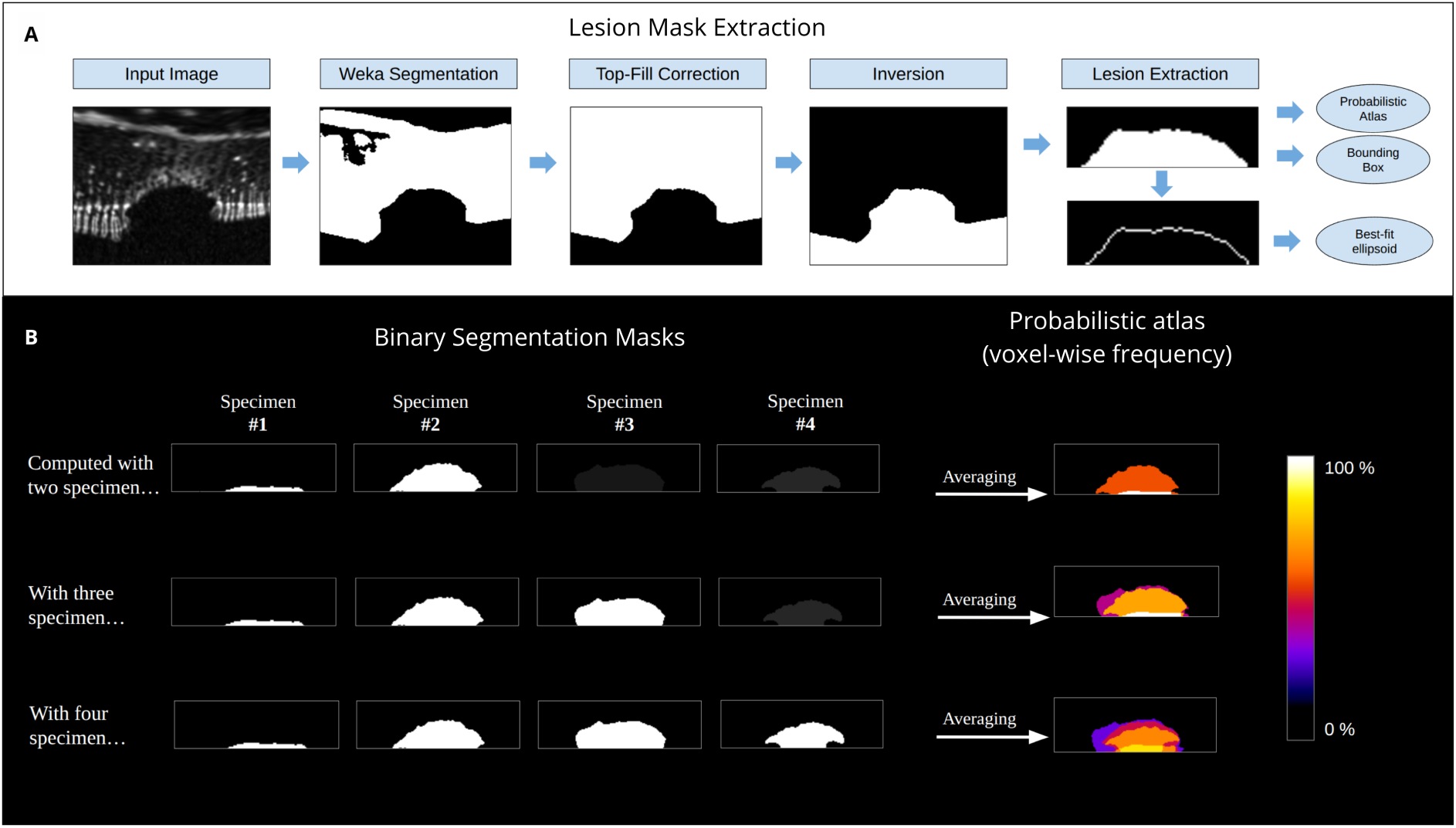
Lesion mask extraction and principle of probabilistic atlas construction. (A) Overview of the lesion segmentation and extraction pipeline, starting from the input MRI slice and proceeding through supervised Weka-based segmentation, top-fill correction, inversion, and final lesion mask extraction. The resulting binary masks were subsequently used to derive geometric descriptors (bounding box and best-fit ellipsoid) and to construct probabilistic atlases. (B) Generation of probabilistic atlases from binary lesion masks. Individual segmentation masks from increasing numbers of specimens were combined by voxel-wise averaging to estimate the frequency of lesion occurrence at each voxel.

### Atlas Construction

#### Mean Images

After spatial alignment, registration, and cylindrical transformation, the images of all specimens in each group were averaged voxel-wise. In the Cartesian space, the average image (Fig 4 (C.1)) appears blurred because the cross-section of the specimen differs in shape and size. This variability increases the pixel-wise standard deviation and smooths the mean signal, hampering the detection of time-lapse spatial trends in the population. After cylindrical transformation, the structures of interest align along a straight axis, resulting in much sharper mean images, with unscattered high intensity area (cambium) and a lower mean variance (Fig 4 (D.1)). The area of high variance we are interested in studying, that appears around the infection zone, is well demonstrated in the cylindrical space (Fig. 4 (D)) highlighting the phenomenon under study, and producing cleaner, more interpretable population-level representations.

Finally, the difference images computed between time points (day 141 and day 1) reveal similar signal patterns in both Cartesian (Fig. 4 (C.3-4)) and cylindrical (Fig. 4 (D.3-4)). Although a stronger contrast enhancement was initially expected in cylindrical space, this was not observed. On the positive side, this result indicates that the cylindrical transformation does not introduce major distortions or artefacts in the signal, and preserves the underlying image content while providing a normalized spatial framework suitable for population-level comparison. Difference images are therefore used here primarily as a qualitative validation of the transformation rather than as an independent source of contrast between using cartesian space and cylindrical space.

#### Probabilistic Atlas

The probabilistic atlases were generated by combining the binary lesion masks and computing, at each voxel, how often the disappearing region (in white) occurred across specimens. The voxel-wise frequency produces a probability map that captures the spatial trends of the lesion geometry within the population. The probability values range from dark purple (0%) to bright yellow (100%), indicating how frequently the lesion appears at each location relative to the inoculation point across the population.

Fig. 5 (B) illustrates this process, showing how individual lesion masks were aggregated into a population-level atlas that highlights regions where lesions overlap consistently across the population (high probability) and regions where specimens diverge (lower probability). Computed with two masks, the atlas shows only three possibilities: 0% (no lesion), 50% (one of the two), or 100% (both). With three masks, the map gains intermediate values, revealing partial overlap (33% or 66%) and full overlap (100%). As more specimens are added, the atlas highlights the population patterns including both common regions (the overlap area) and variable regions of the lesion population. These atlases make it possible to identify group-level trends and to compare how consistently different groups develop the lesion (detailed in the following sections). We applied this to multiple groups to assess the differences: first, Pch group versus Control group (all varieties included), and then one group per variety, including only Pch infected cuttings, see section “Application to monitor and quantify differences between treatments and cultivars”.

### Geometric Features

#### Specimen lesion bounding box

The bounding-box measurements provide a quantitative description of the spatial extent of the lesion in 3D. For each specimen, the box dimensions (height “z”, width “x” and depth “y”) reflect how far the affected region spreads along the different anatomical directions of the cutting. Concisely, larger bounding boxes indicate broader deterioration of internal tissues, while smaller boxes indicate limited/localized progression.

In Fig. 6 (A) the volume in red corresponds to the binary lesion mask, representing all voxels that lost MRI signal due to the fungal colonization. This volume is enclosed by an axis aligned bounding box (in green) which captures the outermost spatial extent of the lesion. Rather than describing lesion volume directly, the bounding box quantifies how far the lesion has spread in each anatomical direction, providing a geometric summary of its lateral, radial, and axial expansion. The x-dimension reflects tangential spread along the cambial region. The y-dimension captures radial penetration toward the trunk centre. The separation of the longitudinal axis into positive and negative z-directions allows independent measurement of apical and basal advancement. Together, these dimensions characterize the 3D extent of the lesion and reveal directional patterns in how the disease progresses through the cutting. However, the bounding-box captures the extremities, making it sensitive to abnormalities that do not correctly represent the actual lesion shape.

**Fig. 6:**
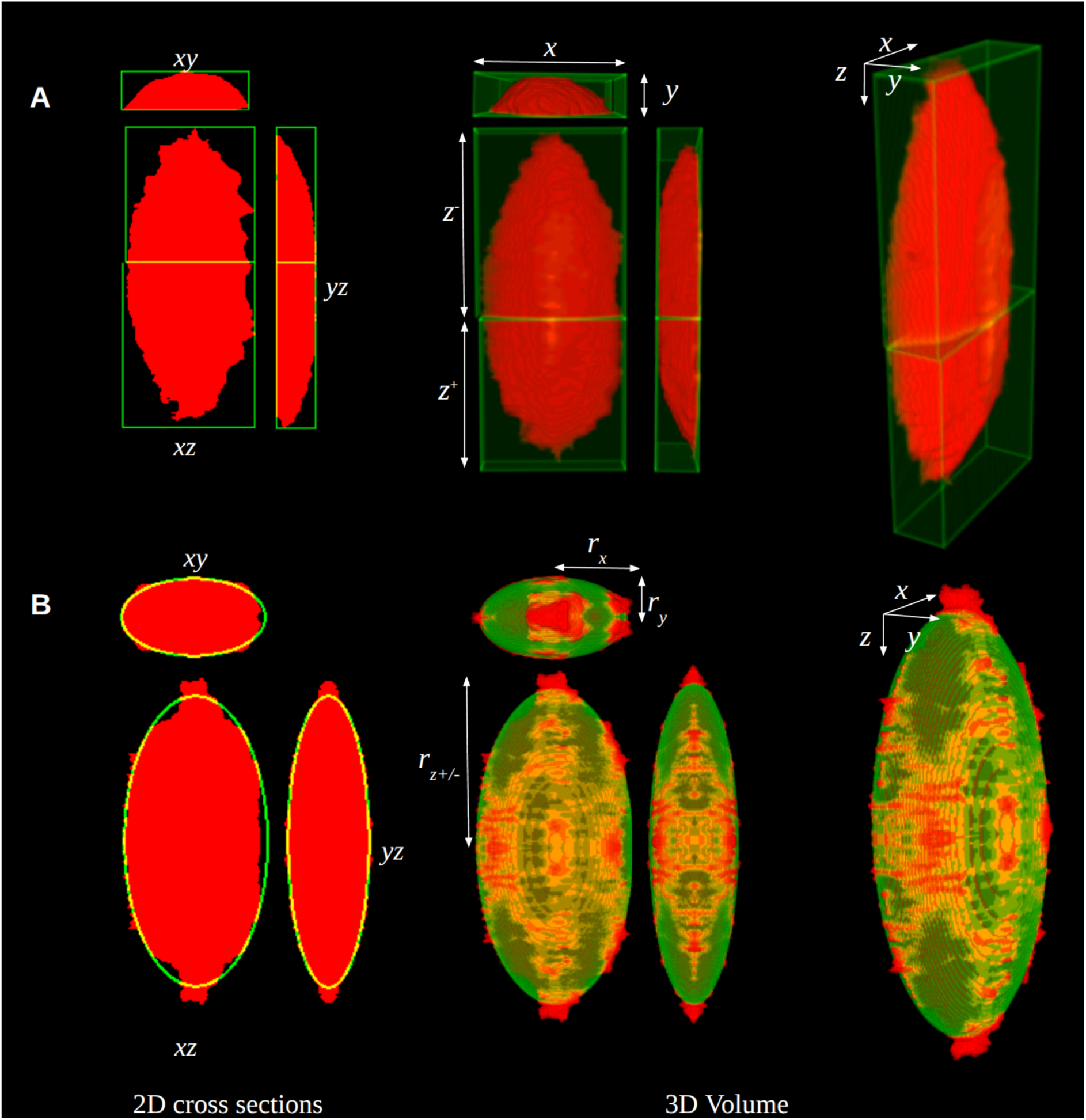
3D visualization of the segmented lesion. (A) Lesion geometry represented using axis-aligned bounding boxes. Orthogonal 2D cross-sections (xy, xz, yz) and a 3D rendering illustrate how the bounding box captures the overall spatial extent of the lesion along the longitudinal (z) and transverse (x, y) axes. (B) Corresponding representation using best-fit ellipsoid modeling. Ellipsoids were fitted to the binary lesion masks to provide a compact and interpretable description of lesion shape and anisotropy.

#### Specimen lesion equivalent ellipsoid

To counter the limitation of bounding boxes, geometric ellipsoids were fitted to the contour of the lesion (Fig. 6 (B)). The average point-wise distance between the actual contour points and the estimated ellipsoids over all specimens is 2.62 *±* 0.91 pixels (compared with the average ellipsoid radii extending between 25 to 100 pixels). This indicates that the fitted ellipsoids closely followed the true lesion contours. Only a small number of samples exhibited high values, reflecting instances where the lesion geometry was more irregular and not well represented by a 3D ellipsoid. Overall, these results confirm that the ellipsoid model provides an adequate geometric representation of the lesions, useful to model their anisotropy with respect to the three anatomical axes.

To ensure that the ellipsoid approximation remained consistent with the lesion geometry, ratios between the ellipsoid radii and the corresponding bounding-box dimensions were also calculated. These ratios provided a simple check that the ellipsoid and the bounding box are not diverging excessively. In our data, the values remained close to 1 across axes, indicating that the fitted ellipsoids captured the lesion shape well and did not underestimate or overinflate its dimensions. This verification step confirms that the fitted ellipsoids are consistent with bounding boxes. However, ellipsoids appeared as a slightly better fit for representing the actual lesion geometry, and were considered more reliable and suitable for subsequent statistical analyses.

### Application to monitor and quantify differences between treatments and cultivars

This section entails the comparison between populations of cuttings, by leveraging the methodology.

#### Pch vs Control

##### Visual investigation of probabilistic atlases

Fig. 7 demonstrates atlases where the colours encode how often a voxel is classified as a lesion across the population, from 0% (dark purple/black) to 100% (bright yellow/white). Only voxels above the cambium central line are considered for the analysis, as voxels below this line correspond to outer tissues (liber, bark), which are not under study.

**Fig. 7:**
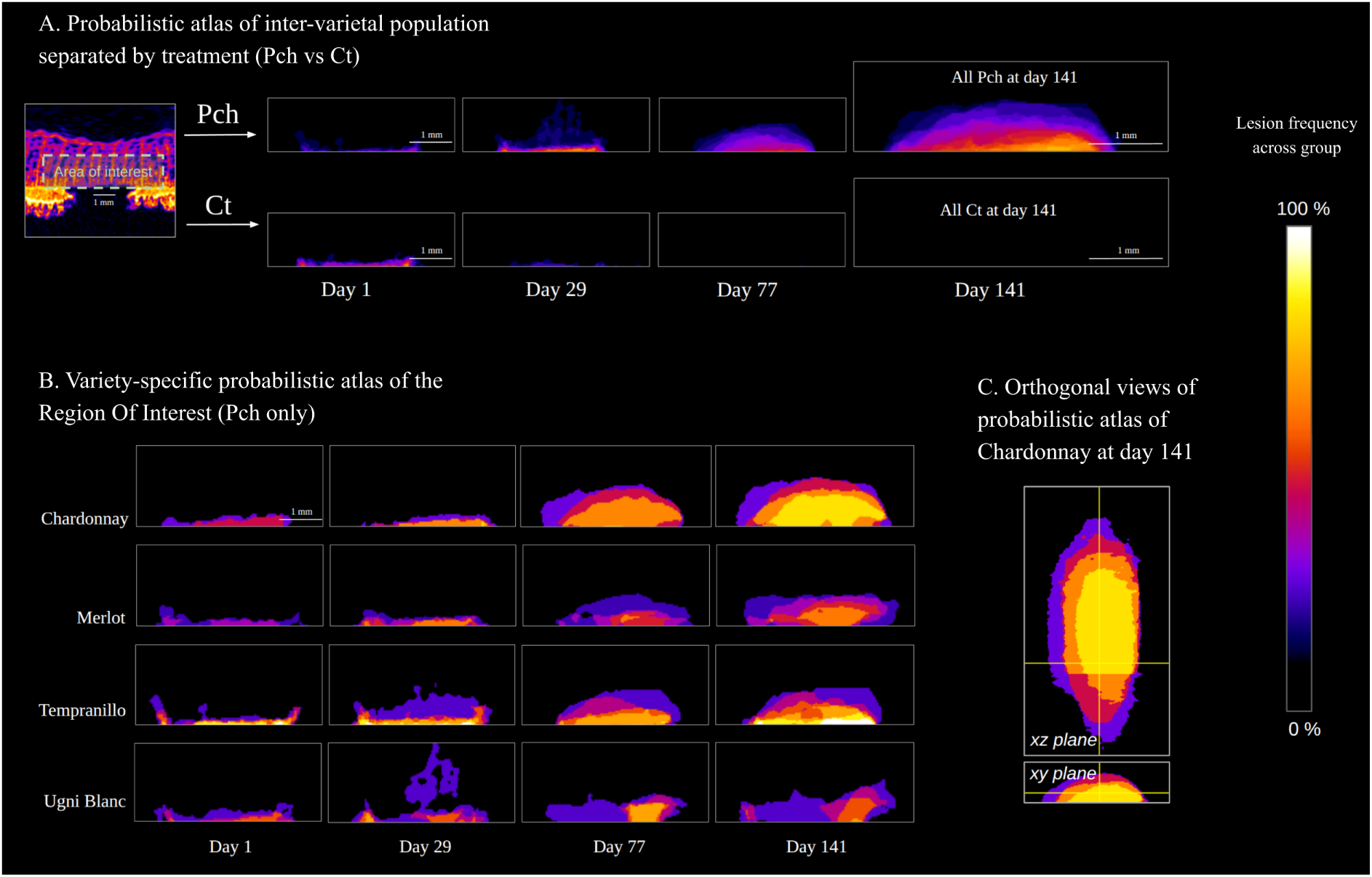
Probabilistic atlases of lesion occurrence across treatments and varieties. Color intensity indicates the voxel-wise frequency of lesion occurrence within each group, highlighting progressive lesion expansion in Pch-infected samples compared with minimal signal in controls. (A) Probabilistic atlases of the region of interest aggregated across all grapevine varieties, shown separately for Pch-infected and Control specimens at successive time points (Days 1, 29, 77, and 141). (B) Variety-specific probabilistic atlases computed from Pch-infected specimens only, illustrating distinct temporal patterns of lesion development among Chardonnay, Merlot, Tempranillo, and Ugni Blanc. (C) Orthogonal views (xy and xz planes) of the probabilistic atlas for Chardonnay at Day 141, illustrating the three-dimensional spatial distribution and extent of lesion occurrence. Scale bars: 1 mm.

The Pch group (infected) shows (Fig. 7 (A)) progressive build-up of high probability necrosis above the threshold (inoculation point) over time, entering the region of interest (inner wood) at the third timestep (day 77) and developing inwards at day 141, with shades of probability indicating a wide distribution of infection depth across the total population of Pch cuttings. Meanwhile, the Control group remains mostly low probability and fades with time, consistent with wound healing and the absence of internal degradation, ultimately exhibiting no deteriorated inner wood at all at the end of experience (day 141).

##### Lesion quantification using ellipsoids

The ellipsoid lesion volumes were computed for each timepoint to quantify how lesion size evolved over the course of the experiment. Each lesion was modeled by fitting separate ellipsoids to the Z⁺ and Z⁻ regions along the stem axis, and their volumes were averaged to produce a single equivalent ellipsoid volume per specimen. This yielded one robust lesion-size estimate per specimen per timepoint, allowing the temporal progression of lesion growth to be compared across cultivars.

Fig. 8 (A) shows that Pch lesions, on average, develop slowly during early infection and then expand rapidly with substantial tissue damage accumulating between day 29 and day 77. In contrast, the absence of a similar volume in Control specimens (segmented volume = 0) further supports the fact that the phenomenon observed was a result of the pathogen action and not linked to the wounding, cultivation setup or imaging protocol.

**Fig. 8:**
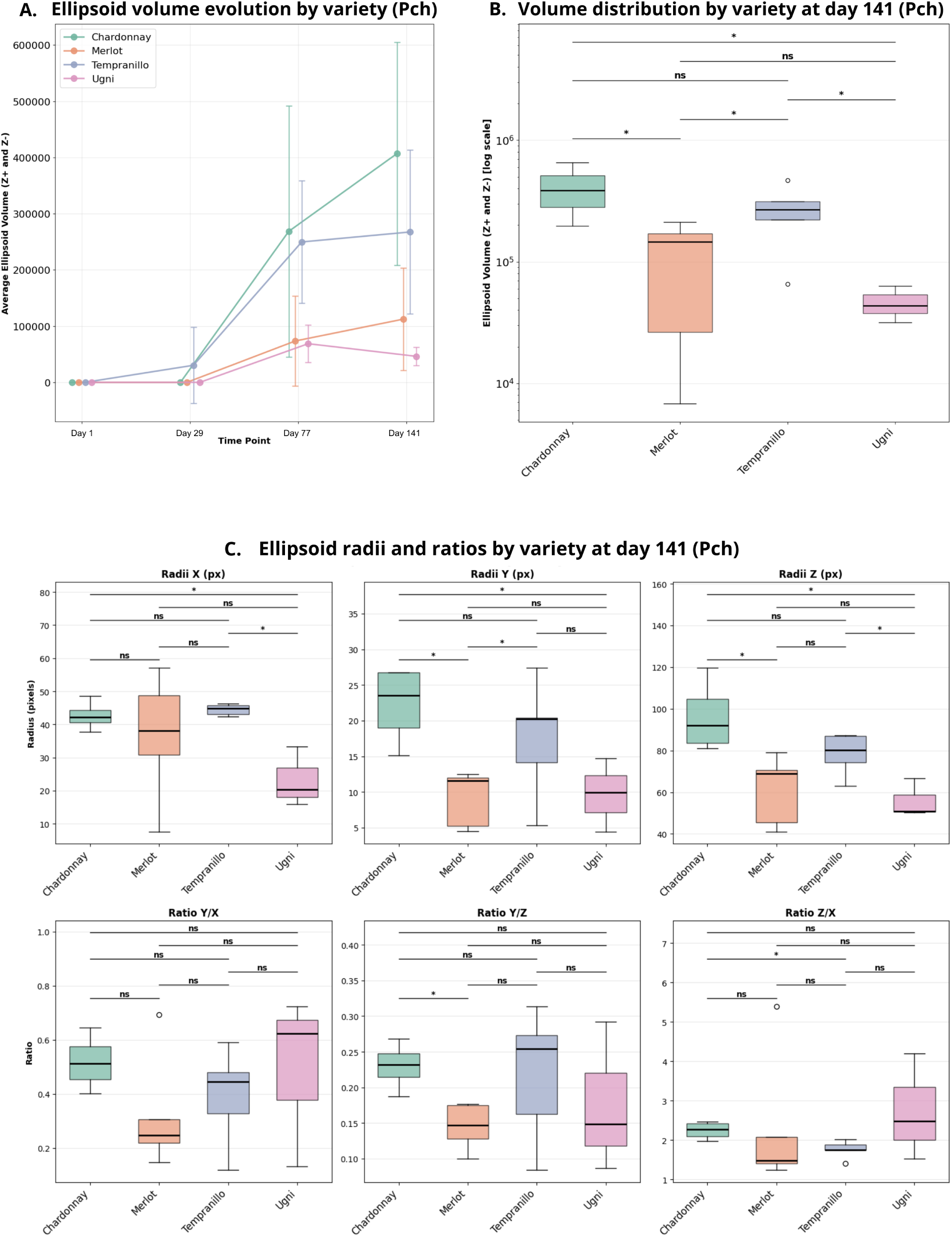
Ellipsoid-based quantification of lesion progression across grapevine varieties under Pch infection. (A) Temporal evolution of average ellipsoid lesion volume for each grapevine variety (Chardonnay, Merlot, Tempranillo, and Ugni) in Pch-infected cuttings. Points represent group means at each time point, with error bars indicating intra-cultivar variability. (B) Distribution of ellipsoid lesion volumes at day 141 for each variety, displayed on a logarithmic scale. Boxplots show median values, interquartile ranges, and individual outliers. Pairwise statistical comparisons between varieties are indicated above the plots (p < 0.05; ns, not significant). (C) Variety-specific distributions of ellipsoid radii along the three anatomical axes (X, Y, and Z) and their corresponding ratios at Day 141, characterizing lesion anisotropy. Boxplots summarize the distribution of each geometric parameter across specimens, with significance annotations indicating pairwise comparisons.

#### Cultivar Differences

##### Visual investigation of probabilistic atlases

We first examined the probabilistic atlas to extract the intra-cultivar variability (Fig. 7 (B)):

- Chardonnay and Tempranillo appeared to be the most consistent varieties, as reflected by probabilistic atlases showing an unscattered area, and the highest ratio between the surface of high probability (common pattern) and the surface of low probability (anomalies).
- On the other hand, Merlot and Ugni blanc exhibit a wide extent of the lesion for a few specimens, while the high probability is limited (in Ugni), or completely absent (in Merlot, the area with maximum probability is at 50%).

We then examined the intra-cultivar trends:

- Chardonnay shows the widest and most consistent high probability region around the inoculation zone (bright yellow, 80% probability, meaning almost all specimens show necrosis in these areas). The orange and red zones extend further towards the pith than in other varieties, indicating a consistently deeper lesion across the cultivar group. In 80 % of the Chardonnay specimen, the lesions reached at least 60% of the biggest lesion depth observed across the population at day 141. Along the x axis, the lesion reached additional tissue from the inoculation area, however this is the case for less than 50 % of the specimen.
- In Tempranillo, all the specimens have developed lesions in the direct proximity of the inoculation area. Along the y axis (depth) the specimen shows a gradient of reactions, the atlas showing a “stairs” pattern with a progressively decreasing lesion probability. Along the x axis (cambium), the lesion reaches no additional tissue outside the inoculation x-axis bounds, making the xylem ray appear as an “invisible wall” in tempranillo.
- Merlot and Ugni Blanc hardly exhibit any trends, as their probabilistic atlases show scattered patterns. The lesions show minimal progress along the y axis, highlighting no common zone reached by all specimens, as a result of variation, with some showing a wide lesion development, and other showing no development at all. Ugni Blanc shows the smallest high probability area among all varieties. Most of the atlas is dominated by purple and magenta, corresponding to low probabilities (10-20%). This colour distribution suggests that most Ugni Blanc specimens experience limited tissue degradation and the lesion rarely expands from the inoculation point. However, both in Merlot and Ugni, some specimens show that once the fungus crosses the cambium surface, the lesion can spread outside the inoculation x-axis bounds, eventually through the xylem rays. This effect is particularly visible in Ugni, with three specimens showing a localized infected area, while in Merlot a single specimen is concerned by this aspect.

##### Lesion quantification using ellipsoids and bounding-boxes

The differences in lesion development progression between cultivars across time was assessed using statistical computation on ellipsoid-based lesion volumes for each cultivar at each timepoint (Fig. 8 (A)). With the exception of Tempranillo, no detectable lesion was observed at day 1 and day 29 in the region of interest, indicating that magnetic resonance imaging during the first 30 days post-inoculation provides no meaningful information on lesion formation. This early absence of signal was consistent across cultivars, and notable lesion development only became visible from day 77 onwards.

The development graph across time exhibits two groups : Chardonnay and Tempranillo, versus Merlot and Ugni, the first group showing big lesions and an ongoing development, while the second group shows smaller lesions, and eventually a stalling lesion development.

Fig. 8 (B) reveal significative differences between cultivars when comparing the development of lesion at the last time-point. The large and rapidly expanding lesions in Chardonnay and Tempranillo indicate high early susceptibility and weak compartmentalization capacity. Merlot demonstrates a more effective limitation of internal damage, while the lesion spread is highly variable across the specimens, like previously seen in the atlases. Ugni Blanc’s minimal lesion volumes suggest enhanced early resistance or more efficient tissue compartmentalization, allowing it to restrict fungal progression even at later infection stages. The coherence between ellipsoid and bounding-box trends confirms that these differences reflect biological variation rather than geometric artifacts.

To test the hypothesis that early lesion development reflects cultivar-specific susceptibility differences, we first formulated the null hypothesis that all varieties exhibit the same lesion volume distribution at the early infection stage. A Kruskal-Wallis test performed on ellipsoid-estimated volumes at Day 141 rejected this null hypothesis (*p* = 0.04) indicating at least one variety differs significantly from the others. This statistically validates the alternate hypothesis that cultivars do not respond uniformly but instead show a distinct level of susceptibility to Pch even at the early stages. The pairwise Kruskal tests (Fig. 8 (B)) show that the cultivar pairs have a significant difference (indicated by their star annotations). Even with the small number of specimens (N=6), significant differences are observed for all comparisons except for Chardonnay vs Tempranillo and Merlot vs Ugni Blanc, where no statistical differences were observed. Overall, the pattern confirms the presence of distinct susceptibility among cultivars during early infection.

It is worth noting that some specimens did not develop any visible lesion during the imaging period and appeared similar to Control plants. No additional information was available to determine whether these vines successfully suppressed the pathogen between Day 1 and Day 29, or whether the inoculation procedure failed for these individuals. These non-responsive specimens were excluded from the statistical analysis to avoid confounding effects, although doing so reduced the sample size and made significance harder to detect. The excluded cases included one Chardonnay, two Merlot, one Tempranillo, and three Ugni Blanc specimens.

Consistent with the volumetric analysis, which show a significant separation of the varieties into two groups with larger (Chardonnay and Tempranillo) versus smaller lesion volumes (Merlot and Ugni Blanc), ellipsoid-derived geometric descriptors (the estimated ellipsoid radii) revealed that this is also reflected along individual dimensions. Chardonnay and Tempranillo exhibited larger extents along r_x_ and r_z_ (Fig. 8 (C) top-left and right) whereas Ugni was significantly smaller along those directions. Differences were also observed in radial lesion depth (rᵧ), with Chardonnay and Tempranillo generally showing deeper penetration compared with Merlot and Ugni Blanc. Merlot showed highly dispersed distributions across all axes, preventing robust statistical discrimination despite occasional large lesions.

When shifting from absolute dimensions to geometric ratios, the grouping became less pronounced. The ratio between r_y_ and r_x_ (Fig. 8 (C) bottom-left) resulted in overlapping distributions across varieties indicating that the radial and depth progression are not discriminant. r _z_/r_x_ (Fig. 8 (C) bottom-right) shows the limitations in the lesion to go through the xylem rays relative to its capability to go through the stem axis, without showing any significant discriminant between varieties. Merlot showed an interesting pattern, particularly in the r_y_/r_z_ ratio (Fig. 8C, bottom-centre), which was approximately two times lower than that of the other cultivars, indicating a lower mean penetration towards the pith relative to the extent along the cambium axis.

## DISCUSSION

This study demonstrates the potential of high-resolution 3D + t MRI to monitor the earliest stages of fungal colonisation in grapevine cuttings, providing functional, non-destructive insight into the tissue functionality during pathogen invasion. Unlike traditional assessments based on external symptoms or destructive sampling, MRI enables continuous observation of internal tissue responses, capturing the onset and progression of lesion formation within days to weeks after inoculation. This early detection capacity is valuable because it reveals physiological changes that precede macroscopic symptoms and may be instrumental for understanding host-pathogen interactions.

A key contribution of this work is the geometric processing framework developed to compare lesions across individuals and timepoints. The combination of landmark alignment (standardizes global pose of the specimens), cylindrical unwrapping (stabilizes anatomical correspondences), population atlases (summarizes populations) and double-symmetry geometric ellipsoid modelling (extracts geometrical descriptors) provides a reliable method for normalising inter-individual variability in stem orientation, anatomy, and lesion shape. Although this framework was developed for *Phaeomoniella chlamydospora* infection in grapevine cuttings, it is generalisable to other host-pathogen systems where internal lesion dynamics needs to be quantified across specimens, especially useful (but not limited to) in combination with magnetic resonance imaging if the pathogen progression induces a loss of host tissue function and a change in water status. The ability to align volumes, extract comparable geometric descriptors, and model lesion morphology systematically opens possibilities for broader comparative pathology studies.

Despite these methodological strengths, several limitations must be acknowledged. While the proof of concept is successful, the sample size remains relatively small, which restricts statistical power, particularly for detecting subtle cultivar differences at early timepoints. Complementary application study on bigger populations with more time points would be interesting to reach a better level of significance. Additionally, the pipeline relies on an assumption of approximate cylindrical symmetry along the longitudinal axis. Although generally valid for young cuttings, this symmetry is not perfect and may introduce minor biases in the ellipsoid fitting process. The cylindrical transformation worked as intended, though its impact was limited by the localized spread of the fungus in this study, and may prove more helpful with longer monitoring experiments or with other pathogens having a wider transversal development. Furthermore, the functioning tissue MRI signal is inherently variable, with varying intensity levels and patterns due to inter-variety or inter-specimen variation of the water content. This has an influence on the lesion/tissue boundaries automatically segmented by the machine learning model, and ultimately impacts volumetric measurements. These factors could be considered when interpreting small differences between individuals or cultivars.

Even with these limitations, the early MRI-derived lesion metrics revealed clear differences among cultivars. In the spatio-temporal patterns, probabilistic atlas highlighted different development patterns, while showing a good reproducibility level of lesion geometry in Chardonnay, an intermediary trend in Tempranillo and big variations in Merlot and Ugni Blanc. The patterns observed suggest a better early compartmentalization by xylem rays in two varieties, while the two others varieties show a capability to limit the inwards progression of the pathogen.

In the study of lesion volume over time, two groups emerged consistently: Chardonnay and Tempranillo showed rapid lesion expansion soon after inoculation, while Merlot and Ugni Blanc displayed slower progression. This separation suggests that early structural degradation captured by MRI reflects meaningful differences in host response. However, this ranking does not fully align with the susceptibility classifications reported in the viticulture literature, where Ugni Blanc is generally described as highly sensitive and Merlot as moderately tolerant. The fact that Tempranillo and Ugni did not follow the expected susceptibility hierarchy is particularly intriguing. This discrepancy suggests that early physiological responses detected by MRI do not necessarily predict long-term disease severity, while short-term water-tissue alterations is the beginning of a process ending in long-term wood necrosis. These observations open important questions about the sequence of processes that connect initial colonisation, host compartmentalisation responses, and the later development of chronic trunk disease symptoms. From the perspective of early indicators, our results show that early lesion formation is indeed informative and can differentiate cultivars by their immediate response to infection, as supported by the significant *p*-value. However, the divergence between early MRI-based susceptibility patterns and long-term vineyard observations emphasises the need for extended longitudinal studies. Continuous monitoring over months or seasons would help determine how early physiological signatures translate into later structural degradation and whether the initial MRI signal loss predicts eventual disease outcomes.

Going forward, several developments could strengthen the approach. Higher temporal resolution (smaller timestep) imaging could capture the very early transition between functional tissue and altered one, with a detectable signal loss. Multi-zone analysis separating conductive tissues (xylem vessels) from structural wood (xylem rays) may provide a more nuanced understanding of how the fungus perturbs hydraulic function. Finally, integrating geometric lesion metrics with mechanistic modelling of pathogen growth and host responses could help explain how different cultivars regulate or fail to regulate infection spread.

Our work can have practical implications for grapevine health management such as early-stage assessment of internal tissue degradation, where rapid identification of sensitive genotypes can help with planting decisions and guide disease prevention strategies. This framework is not restricted to the four cultivars or the single pathogen studied here. It can be readily applied to other grapevine varieties and trunk disease fungi for further comparative studies. In the current phytosanitary context, where the single effective treatment product (arsenite) is restricted in the EU, growers have limited options beyond physically removing infected wood. This work highlights the needs for tools that can detect internal infections before they require destructive interventions, positioning MRI as a non-invasive and dynamic approach. In addition, the quantitative nature of the pipeline makes it a promising benchmark for evaluating the efficacy of new phytosanitary treatments being deployed.

Overall, this work demonstrates that MRI 3D + t, combined with geometric lesion modelling, offers a powerful framework for non-destructive early-stage monitoring of grapevine trunk disease. It reveals cultivar-specific early responses, uncovers unexpected discrepancies with long-term susceptibility classifications, and provides methodological foundations for future studies linking early physiological changes to chronic disease development.

## ACKNOWLEDGEMENTS

The authors thank all colleagues and technical staff who contributed to plant cultivation, sample preparation, MRI acquisitions, and experimental support. We also acknowledge valuable discussions and logistical assistance that helped facilitate this study. Special thanks to Denis Cornet, Alain Audebert, and Michel Ghanem for their contributions to funding the computing servers, and to #DigitAg “Digital Agriculture Convergence Lab” (https://www.hdigitag.fr) for its supportive community.

## Author contributions

GP: Conceptualization, Methodology, Software, Validation, Formal analysis, Investigation, Data curation, Visualization, Writing - original draft, Writing - review & editing.

MC: Investigation, Resources, Data curation, Methodology, Writing - review & editing.

CGB: Investigation, Resources, Data curation, Methodology, Writing - review & editing.

LLC: Conceptualization, Supervision, Project administration, Funding acquisition, Writing - review & editing.

JLV: Conceptualization, Resources, Supervision, Project administration, Funding acquisition, Writing - review & editing.

CM: Conceptualization, Methodology, Investigation, Resources, Supervision, Project administration, Funding acquisition, Writing - review & editing.

RF: Conceptualization, Methodology, Software, Validation, Formal analysis, Visualization, Supervision, Funding acquisition, Writing - original draft, Writing - review & editing.

## DATA AVAILABILITY

The datasets generated and analyzed in this study include approximately 160 GB of raw 3D magnetic resonance images and 1.4 TB of processed data products. Due to the large volume of these files and current storage and transfer constraints, the data are available from the corresponding author upon reasonable request. The processing pipeline (including scripts and parameters required to reproduce the processed outputs from the raw data) is available at https://doi.org/10.5281/zenodo.17944369.

## SUPPLEMENTARY MATERIALS

### Note S1

**Fig. S1:**
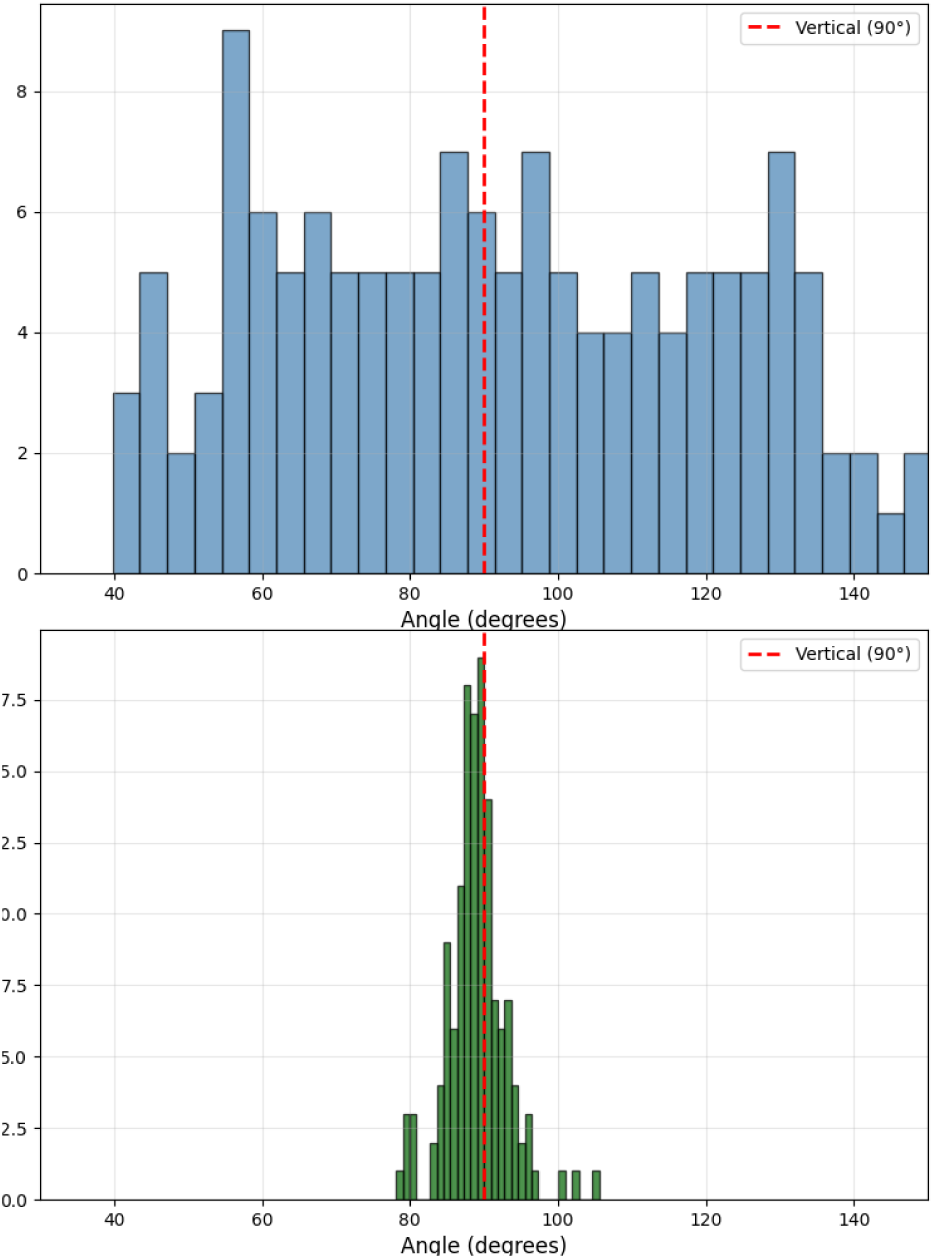
Xylem rays angle distribution - before transformation (Cartesian)

**Fig. S2:**
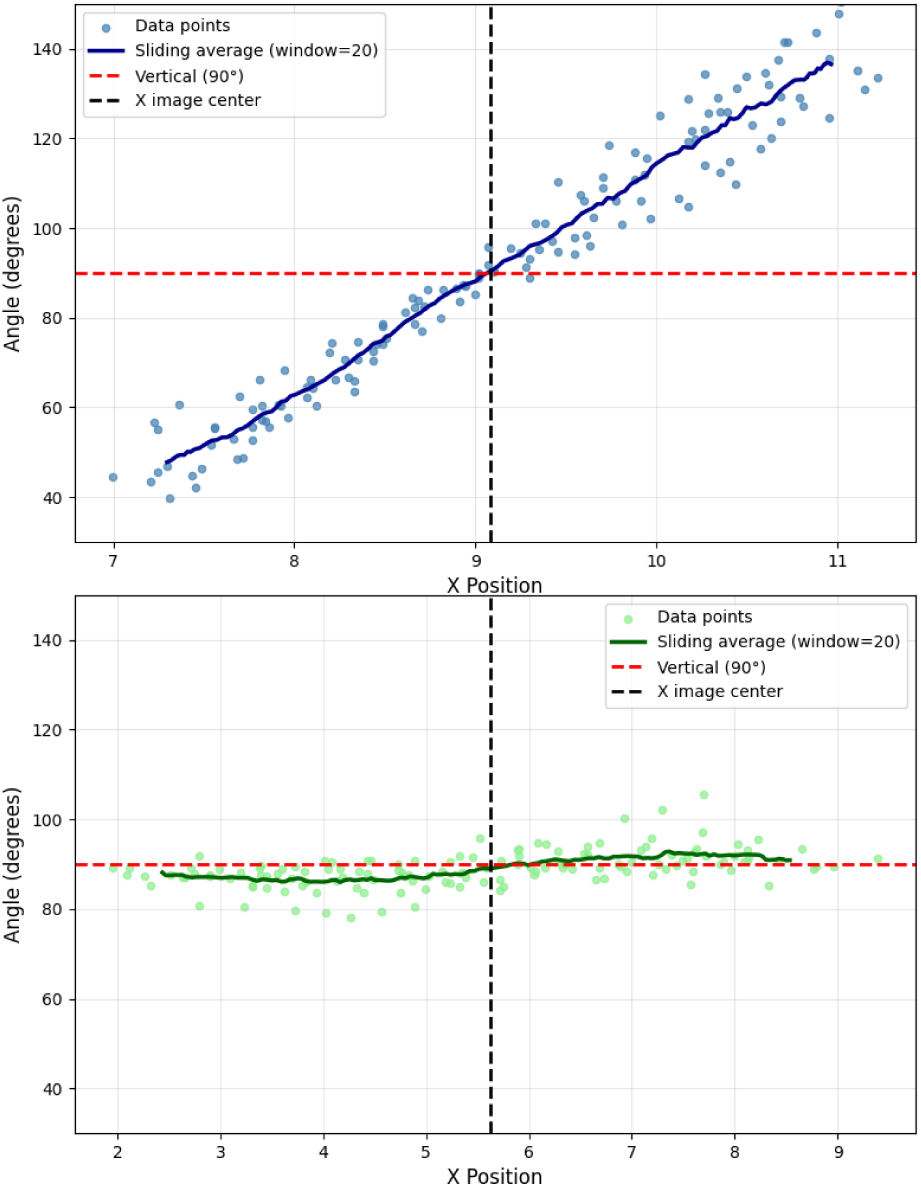
Angle vs X Position - after transformation (polar)

